# Functional Connectome Fingerprinting Using Shallow Feedforward Neural Networks

**DOI:** 10.1101/2020.10.19.346189

**Authors:** Gokce Sarar, Bhaskar Rao, Thomas Liu

## Abstract

Although individual subjects can be identified with high accuracy using correlation matrices computed from resting-state functional magnetic resonance imaging (rsfMRI) data, the performance significantly degrades as the scan duration is decreased. Recurrent neural networks can achieve high accuracy with short duration (72s) data segments but are designed to use temporal features not present in the correlation matrices. Here we show that shallow feedforward neural networks that rely solely on the information in rsfMRI correlation matrices can achieve state-of-the-art identification accuracies (≥ 99.5%) with data segments as short as 20s and across a range of input data-size combinations when the total number of data points (# regions × # time points) is on the order of 10,000.

---

Functional connectome fingerprinting based on the similarity of correlation coefficient matrices computed from rsfMRI data can identify individuals with high accuracy (> 98%) using long duration (> 12 minute) scans but considerably lower accuracy (≈ 68%) is obtained when the data duration is decreased to 72s (1). Recurrent neural networks (RNN) can achieve high accuracy (98.5%) with short duration (72s) data, presumably reflecting their ability to capture both spatial and temporal features (2, 3). However, it has been shown that high RNN performance can be achieved even when the temporal order of the fMRI data is permuted (4), suggesting that the temporal features are not critical for identification. Here we introduce two shallow feedforward neural networks that can achieve high identification accuracy without the need for recurrent connections. Furthermore, we use these networks to estimate the minimum size of the data needed to robustly identify subjects with high mean accuracy (≥ 99.5%) from short segments of rsfMRI data. Since identification accuracy reflects the ability to effectively extract information from functional connectomes, additional insight into the methods and minimum data sizes that achieve high performance can guide the development of extended approaches to detect other differences in functional connectivity, such as disease-related changes.

The two networks considered are shown in Figure 1A and 1B. The input to the correlation neural network (corrNN) consists of the upper triangular elements of the correlation coefficient matrix *C* estimated from a data matrix *X* consisting of z-normalized time series (of length *N*) from *M* regions of interest (ROI). For identification of *L* subjects, the network structure consists of a fully connected classification layer with *L* units, a batch normalization layer, and a softmax layer. The norm-based neural network (normNN) uses the z-normalized data *X* as the input. The first stage is a fully connected layer that projects the data onto *K* hidden units using the *M* × *K* weight matrix *W* to form the *N* × *K* intermediate matrix *Y* = *XW*. In the second stage, the *L*_2_ norm across the time-dimension (i.e. across each column of *Y*) is computed for each hidden unit to form a summary measure of similarity over the collection of *N* time points. The resulting vector 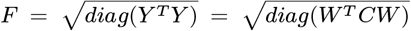 is comprised of *K* features extracted from the correlation matrix *C*. The *K*th feature is proportional to the variance in the direction of the *K*th column vector of *W*. If these vectors are randomly oriented and constrained to be unit norm, then the features represent a random sampling of the “peanut” shaped surface of directional variances (5). The subsequent stages in the network are: a batch normalization layer, a fully connected classification layer with *L* hidden units, a second batch normalization layer, and a softmax layer.

**Fig. 1.**
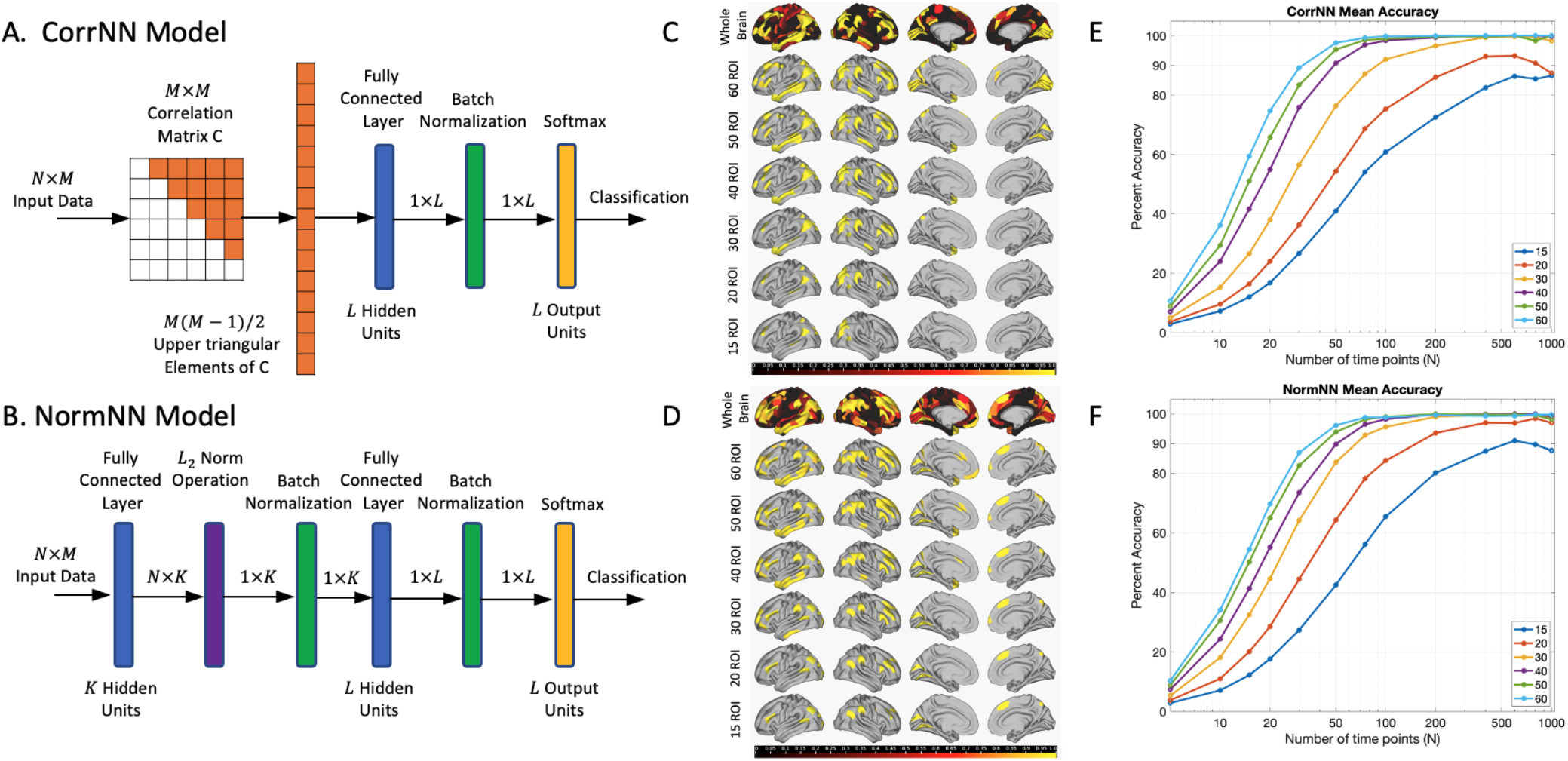
(A,B) CorrNN and NormNN model structures. (C,D) Top rows: Maps showing the relative importance of the ROIs for identification accuracy with maximum importance of 1.0 indicated in yellow. The remaining rows are thresholded to show the locations of the top 15 to 60 ROIs. (E,F) Mean identification accuracies as a function of the number of time points and ROIs.

## Results

We assessed the performance of the two networks using data from the Human Connectome Project (HCP) (6). Two rsfMRI scans acquired on Day 1 were used for training, while the two scans from Day 2 were used for validation and testing.

For *M* = 379 ROIs, *N* = 100 time points (72s duration) per segment, and *K* = 256 hidden units, the mean classification accuracies of the corrNN and normNN models were 99.8% and 99.6%, respectively, for an initial set of 100 subjects, and 100.0% and 99.7% for a second independent set of 100 subjects. These accuracies are higher than those reported (94.3% to 98.5%) for RNN models (2, 3). For comparison, the mean classification accuracy using the similarity of the correlation coefficients was 79.4% for 100 time points per segment, which is higher than the 68% mean accuracy reported in (1) using data from a different dataset.

We used a greedy search algorithm to assess the relative importance of the ROIs with respect to model accuracy. Importance maps are shown in the top rows of Figures 1C and 1D, respectively, with the subsequent rows thresholded to highlight the top 15 to 60 ROIs. When considering the top 60 ROIs, the highest number of ROIs are found in region 22 (dorsolateral prefrontal cortex) followed by regions 17 (inferior parietal cortex), 14 (lateral temporal cortex), 16 (superior parietal cortex; for CorrNN), 21 (inferior frontal Cortex), and 3 (dorsal stream visual cortex), where brain regions are as defined in (7).

We used the top ROIs to evaluate CorrNN and NormNN performance with 15 to 60 ROIs and 5 to 1000 time points, as shown in Figures 1E and 1F, respectively. As the number of ROIs decreases, the number of time points needed to achieve higher accuracy increases. Defining 99.5% as the threshold for high mean accuracy, we observed that this threshold is surpassed with as few as *M* = 60 ROIs and *N* = 100 time points for CorrNN and 40 ROIs and 200 time points for NormNN, corresponding to *M* × *N* = 6000 or 8000 total data points, respectively.

To further explore the dependence on number of ROIs and time points, we considered combinations (*M, N*) where the total number of data points was constrained to be equal to or close to either 6000 or 10,000 (see Figure 2 caption). For CorrNN, high mean accuracies are obtained for two of the combinations (dark red squares) with 6000 data points and all five of the combinations (dark red diamonds) with 10,000 points, respectively.

**Fig. 2.**
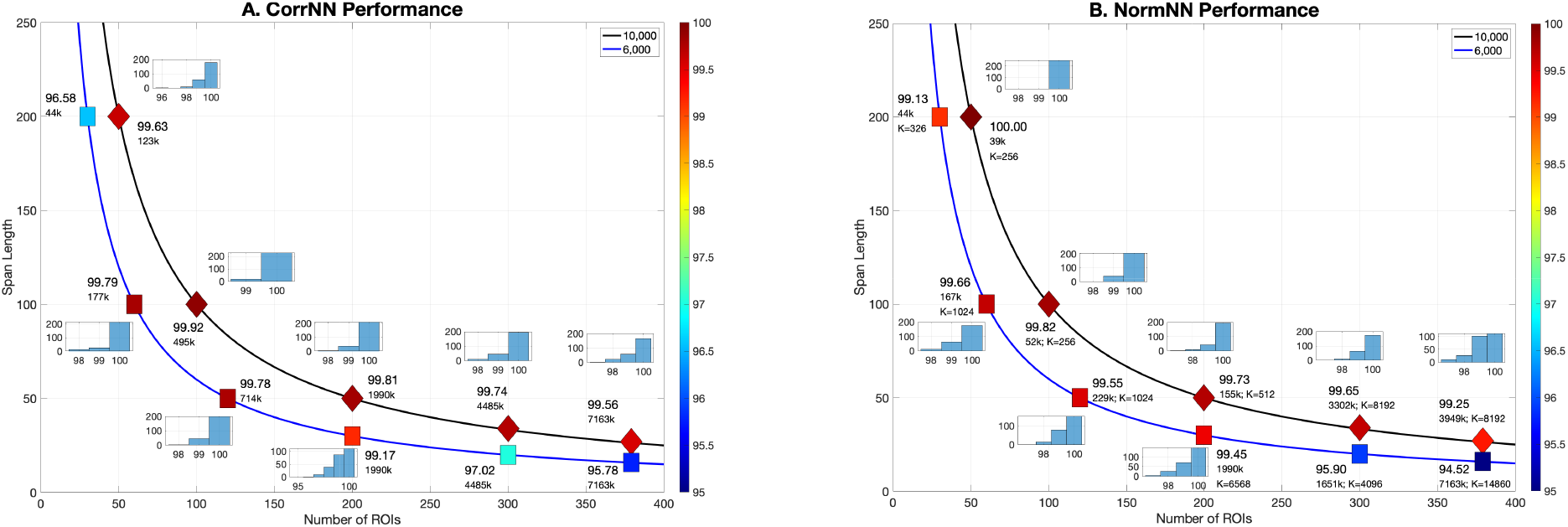
(A,B) CorrNN and NormNN identification accuracies for combinations (*M, N*) of numbers of ROIs (*M*) and span lengths (*N*) that are constrained to have either 6000 or 10,000 data points, with the exception of the combinations (379,16), (300, 34), and (379, 27), which have (5064, 10,200, and 10,233 data points, respectively. Mean accuracies are indicated by labels and color scale. The numbers of model parameters (in thousands) for CorrNN (0.5*L* (*M*^2^ — *M* + 6)) and NormNN (*K* (*M* + *L* + 3) + 3*L*) are also listed (with *L* = 100), as are the numbers of hidden units (*K*) for NormNN combinations. For combinations where CorrNN mean accuracy is greater than 99%, auto-scaled histograms show the distribution of identification accuracies obtained over 250 test trials per combination.

For NormNN, the number of parameters exhibits a linear dependence on the number of ROIs (*M*) as compared to the quadratic dependence for CorrNN (see Figure 2 caption). To better compare the models, we increased *K* by powers of 2 up to the value *K_eq_* = 0.5L (*M*^2^ — *M*) /(*M* + *L* + 3) for which the numbers of NormNN and CorrNN parameters were equivalent, while also including *K_eq_* as one of the possible options. In figure 2b, we show NormNN accuracies obtained for either (1) the minimum value of *K* ≥ 256 that surpassed the 99.5% threshold or (2) the value *K* ≤ *K_eq_* that achieved the highest accuracy when the threshold was not met. High mean accuracies were obtained for two and four of the combinations with 6000 and 10,000 data points, respectively.

As shown by the histograms, the high mean CorrNN and NormNN accuracies correspond to robust identification performance with the majority of the trials demonstrating 100 percent prediction accuracy. These accuracies were obtained with global signal regression (GSR), and were significantly greater than those obtained without GSR for both CorrNN (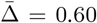; *t*_10_ = 5.00; *p* = 0.0005) and NormNN (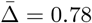; *t*_10_ = 3.93;*p* = 0.0028) where 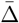 denotes the mean difference in accuracy. Without GSR, only two of the CorrNN combinations and two of the NormNN combinations exhibited accuracies greater than 99.5%.

Using the ROIs determined from the first 100 subjects we evaluated performance on the 2nd set of 100 subjects for the combinations denoted in Figure 2. High mean CorrNN accuracies (≥ 99.5%) were maintained for both of the previously identified high performance combinations with 6000 points and four of the combinations with 10, 000 points, with the remaining combination (379, 27) exhibiting slightly lower accuracy (99.28%) for the 2nd dataset. Thus, the same set of ROIs can offer comparable and high levels of performance across independent datasets.

For both sets of subjects, the mean numbers of CorrNN prediction errors were not significantly correlated (across subjects) with mean framewise displacement (FD) measures of subject motion (|*r*| < .05; *p* > 0.64). Correlations were higher but did not reach significance when using a filtered version of the FD measures (see SI: Extended Methods) with values of *r* = 0.19 (*p* = 0.06) and *r* = 0.13 (*p* = 0.19) for the first and second subject groups, respectively. When viewed within the context of the high CorrNN accuracies that can be achieved, these results suggest that any effects of subject motion on performance are fairly weak.

For NormNN we find that the first layer trained weights are randomly distributed so that the features after the *L*_2_ norm operation represent an approximately uniform sampling of the directional variance surface of *C*. Indeed, high performance can also be achieved by replacing the first layer with a set of random Gaussian weights. The generalizability of the features across datasets exhibits a dependence on the number of units *K*. For example, when using first layer weights trained using the first set of subjects, performance for the combination (100,100) with *K* = 256 drops from 99.82% for the first 100 subjects to 98.79% for the second 100 subjects. Increasing to 1024 units with weights trained using the first set yields accuracies of 99.91% and 99.63% for the first and second sets, respectively. Comparable accuracy levels (99.81% and 99.65%) are obtained when using random weights for the first layer. Thus, generalizability of the NormNN features increases when there is a higher number of features to characterize the directional variance.

## Discussion

We have shown that shallow feedforward models can identify subjects based solely on information in rsfMRI correlation matrices, with CorrNN directly using the correlation coefficients as features while NormNN uses features related to the directional variance surface. The performance levels achieved are state-of-the-art, with high (≥ 99.5%) mean identification accuracies robustly obtained with 6000 to 10,000 data points. For comparison, the convolutional RNN presented in (3) achieved 98.5% accuracy with 23, 600 data points.

Consistent with prior observations (1), high performance can be achieved when using a subset of the ROIs, including those located in frontoparietal and lateral temporal regions. The same set of ROIs can be used to achieve high performance across independent datasets, suggesting that the predictive value of inter-subject variability in the functional boundaries and connectivity of these regions generalizes across datasets.

While combinations with span lengths as short as 27 points (19.5s; CorrNN (379, 27)) can offer high performance, they require a large number of model parameters. In contrast, combinations with fewer ROIs but increased span lengths (e.g. (100, 100)) achieve high performance with one to two orders of magnitude fewer parameters. For NormNN the number of trainable parameters can be further decreased through the use of random weights in the first layer.

As in prior studies (1–3), the current study utilized the HCP dataset, in which the data were acquired on two consecutive days. Although substantial variations in functional connectivity can occur on short time scales (i.e. minutes to hours) due to factors such as temporal fluctuations in subject arousal and vigilance (8, 9), our results indicate that high performance can be obtained over a 1-day interval even in the presence of these factors. Future large-scale studies with a greater time interval between training and test data will be needed to assess whether high identification accuracy can be obtained over longer intervals (i.e. weeks to years).

The effectiveness of the feedforward networks for distinguishing individuals with relatively little data suggests that similar future approaches may have the potential to more fully utilize the information contained in rsfMRI data to better identify disease-related differences.

## Materials and Methods

Details of the methods are provided in Supplementary Information: Extended Methods. Data and analysis code are available at: https://bitbucket.org/ttliu07/fc_fingerprint_nn

## ACKNOWLEDGMENTS

This work was supported in part by NIH grant R21MH112155. We thank Eric Wong, Garrison Cottrell, Jiawei Ren, and Shili Wang for their assistance and helpful comments.

## Supplementary Information Text

### Extended Methods

#### Data

The data used in the study are from the Human Connectome Project (HCP) (1). The first set of 100 Subjects are from the 100 Unrelated Subjects subset of the HCP 1200 dataset, while the second set of 100 subjects included 100 additional subjects from the S900 subjects release of the HCP 1200 dataset. For each subject, the data consisted of 4 resting state scans from 2 separate sessions (2 scans per session), where the sessions were performed on two consecutive days and each scan is 14.4 minutes long with 1200 time points and a repetition time of 0.72 seconds.

#### Preprocessing

Details of the preprocessing performed by the HCP are described in (2–4) and included realignment to correct for subject motion, linear detrending, and ICA based denoising. The data were registered to a common cortical surface using a multimodal registration method (MSM-All) (5). We divided the data into 379 regions of interest (ROI) consisting of 360 ROIs in the cortex as defined in (6) and 19 subcortical ROIs as defined in the group average parcellation provided by the HCP (1). The time series were averaged within each ROI and global signal regression (GSR) was applied to the ROI time series (7). Supplementary analyses were also performed on the data without application of GSR.

#### Training, Validation, and Testing

We used the ROI-averaged data from day one as the training set (2 scans; 1200 pts per scan) and data from day two as validation and test sets (1 scan each). Overlapping data segments were used with shifts of 1 point between segments for training and validation and 250 randomly chosen initial starting points for testing.

The full connected layers were initialized with the Glorot uniform initializer (8). We used the Adam optimizer with learning rate, *β*_1_, and *β*_1_ values of 0.001, 0.9, and 0.999, respectively. We used a batch size of 64 and monitored the validation loss every 600 steps with patience set to 30 monitoring steps and the learning rate annealed by halving it every 100 monitoring points.

The performance of the models was initially assessed using *M* = 379 ROIs and *N* = 100 time points. Using the trained model weights, we assessed the relative importance of the ROIs by first looping over all 379 ROIs, independently zeroing out the data from each ROI, and finding the ROI for which the model retained its maximum accuracy. The identified ROI data were eliminated (i.e. set to zero) for the remainder of the process and the search was repeated over the remaining 378 ROIs to select the next ROI for elimination. This process was continued until only one ROI remained. The importance score of each ROI was 1 — *M_c_* where *M_c_* denotes the model accuracy just prior to ROI elimination.

We then examined performance (shown in Figure 1) across a range of ROI numbers (*M* = [15, 20, 30, 40, 50, 60]; with the top *M* ROIs selected based on importance scores) and durations (*N* = [5,10,15, 20, 30, 50, 75,100, 200,400, 600, 800,1000] time points), retraining the models for each combination (*M,N*) with 1 point shifts for both training and validation. Testing was performed with 250 randomly chosen initial starting points with the exception of 200 sequential points for *N* = 1000. Performance using additional parameter combinations (detailed in Figure 2 and associated text) was also evaluated.

We used the approach of (10) to identify subjects based on similarity of the correlation coefficient matrices. Target matrices were calculated using all the data from day one, whereas test matrices were calculated using data segments from day two. Identification was performed by computing the spatial correlation between the test and target matrices and then matching (with replacement) each test matrix to the most highly correlated target matrix.

#### Subject Motion and Identification Accuracy

For each resting-state scan, the frame-wise displacement (FD) was computed from the time series of the six motion parameters (3 translation parameters and 3 rotation parameters) calculated from the image realignment performed during preprocessing (2, 11). As respiration-related image artifacts can contaminate motion parameter estimates for fMRI data, especially when the data are acquired with short repetition times such as those used in the HCP (12, 13), we followed the approach of (14) and lowpass filtered (0.1 Hz) the motion parameters and also computed filtered FD (fFD) estimates. The mean FD and fFD values were then calculated for each subject by averaging over time points and scans. The two subject groups did not differ in either mean FD (*p* = 0.35) or fFD values (*p* = 0.59), with group FD means (± standard deviation) of 0.167 ± 0.062 mm and 0.172 ± 0.070 mm for the first 100 and second 100 subjects, respectively, and corresponding fFD means of 0.021 ± 0.013 mm and 0.021 ± 0.014 mm.

For each combination ((*M,N*) and number *K* of first layer units for NormNN) evaluated in Figure 2, we recorded the number of prediction errors per subject and then computed each subject’s mean number of prediction errors by averaging across the combinations. For each of the subject groups (first or second 100 subjects), the mean numbers of prediction errors were then correlated with the mean FD and mFD values across all subjects in the group.

## Notes

### Competing Interest Statement

The authors have declared no competing interest.

